# A semi-quantitative, rapid, point of care SARS-CoV-2 serologic assay predicts neutralizing antibody levels

**DOI:** 10.1101/2023.05.30.542314

**Authors:** Alena J. Markmann, D. Ryan Bhowmik, Baowei Jiang, Michael Van Hoy, Frank Wang, Yixuan J. Hou, Ralph S. Baric, Aravinda M. de Silva, Luther A. Bartelt

**Affiliations:** Department of Medicine, Division of Infectious Diseases, University of North Carolina School of Medicine, Chapel Hill NC 27599, USA; Department of Microbiology and Immunology, University of North Carolina School of Medicine, Chapel Hill NC 27599, USA; BioMedomics Inc. Morrisville, NC 27560, USA; Department of Epidemiology, School of Public Health, University of North Carolina, Chapel Hill, NC, USA; Moderna Therapeutics Inc., Cambridge, MA, USA

## Abstract

The ongoing COVID-19 pandemic has caused millions of deaths and the continued emergence of new variants suggests continued circulation in the human population. In the current time of vaccine availability and new therapeutic development, including antibody-based therapies, many questions about long-term immunity and protection remain uncertain. Identification of protective antibodies in individuals is often done using highly specialized and challenging assays such as functional neutralizing assays, which are not available in the clinical setting. Therefore, there is a great need for the development of rapid, clinically available assays that correlate with neutralizing antibody assays to identify individuals who may benefit from additional vaccination or specific COVID-19 therapies. In this report, we apply a novel semi-quantitative method to an established lateral flow assay (sqLFA) and analyze its ability to detect the presence functional neutralizing antibodies from the serum of COVID-19 recovered individuals. We found that the sqLFA has a strong positive correlation with neutralizing antibody levels. At lower assay cutoffs, the sqLFA is a highly sensitive assay to identify the presence of a range of neutralizing antibody levels. At higher cutoffs, it can detect higher levels of neutralizing antibody with high specificity. This sqLFA can be used both as a screening tool to identify individuals with any level of neutralizing antibody to severe acute respiratory syndrome coronavirus 2 (SARS-CoV-2), or as a more specific tool to identify those with high neutralizing antibody levels who may not benefit from antibody-based therapies or further vaccination.

## Introduction

There is currently a need for rapid quantitative antibody assays that assess an individual’s immune response or the lack thereof to SARS-CoV-2. Rapid quantitative antibody assays can be used to identify individuals who may benefit from repeated vaccination given recent studies showing strong positive correlations between breakthrough infections and neutralizing antibody titers[1], antibody-based therapeutics, as a diagnostic aid for individuals with negative molecular testing, and to identify those who qualify to donate antibody-based therapies. In order for serologic assays to inform clinical decisions and public health interventions, antibody-based correlates of immune protection and their duration need to be established, and quantitative antibody assays need to be developed and validated against more rigorous functional antibody assays[2].

Binding and functional neutralizing antibody responses to natural SARS-CoV-2 infection as well as vaccination are highly variable in both duration and titer[1, 3-6]. Though the humoral immune response is only one component of the immune response to SARS-CoV-2, it is important to understand this variability and how it effects durability and protection against future infection. Furthermore, clinical trials evaluating antibody-based therapies for the disease caused by SARS-CoV-2 or COVID-19 have identified a clear benefit for participants enrolled prior to antibody seroconversion[7]. In this study, we use a large cohort of individuals naturally infected by SARS-CoV-2, pre-vaccination and at convalescence, to evaluate the correlation of a linically validated serologic lateral flow assay[8] with a functional antibody neutralization assay[9] and show that it can be used to identify individuals with detectable neutralizing antibody levels.

## Materials and Methods

This study was conducted under Good Clinical Research Practices and compliant with institutional IRB oversight approved by the UNC IRB (#20-1141), consent was obtained from all participants. 268 convalescent SARS-CoV-2 plasma samples from natural pre-delta variant infection prior to any vaccination availability were used for this investigation. These samples were collected at a median of 62 days post PCR diagnosis (n = 215) or symptom onset (n = 53),whichever came first, with a range of 12-337 days. For the BioMedomics qLFA, which has been separately validated[8],10 uL of serum or plasma was pipetted onto a SARS-CoV-2 receptor binding domain (RBD) IgG test strip, followed by three drops of buffer solution provided with the kit per manufacturer instructions. The LFA has been previously validated for whole blood as well as different types of venous samples including plasma[8]. Each strip was developed for 13-15 minutes at room temperature with standard lighting conditions. The strip was then inserted into a prototype RI detector (see Supplemental Figure 1), which displayed a qualitative result (Positive/Negative/Intermediate) and a quantitative result in the form of reflective intensity (RI) of gold particles on the LFA strip. The RI linear range ranged from 0 to 3000. Values were considered to be positive according to manufacturer protocol: RI >80, intermediate: RI 50-80, and negative: RI <50. To determine linearity of the RI detector, human IgG antibody was diluted in human serum at different concentrations. Each sample was then tested on the previously validated Biomedomics IgG RBD LFA and read by the detector. The data was then used to create a calibration curve for the detector.

Results from the sqLFA were then compared to an in-house RBD IgG enzyme-linked immunosorbent assay (ELISA)[5, 10] and a live virus luciferase reporter-based functional neutralization assay whose readout is 50% neutralization of viral infection (NT50)[9]. These assays were all performed as previously described[5].

### Statistics

All statistical analyses and graphs were generated using GraphPad Prism 9.2.0 for Windows[11]. A non-parametric Spearman correlation coefficient was calculated using GraphPad Prism to compare the BioMedomics sqLFA RI to the NT50 titer and in-house RBD IgG ELISA quantitative end-point titer. All tests were two-tailed and a p-value less than 0.05% was considered statistically significant. A receiver operating characteristics curve (ROC) was conducted to determine the sensitivity and specificity of the BioMedomics qLFA in detection of functional neutralizing antibody levels.

## Results

### sqLFA performance

We found that the sqLFA RI readout for the samples tested ranged from 0-2169, and were positively correlated with NT50 titers (See Supplemental Figure 2A), with a Spearman correlation coefficient (r_s_) of 0.70 (p < 0.0001, n = 268). We also obtained the Spearman correlation coefficient (r_s_) for samples that were 14-59 days, ≥ 60 days or ≥ 90 days post diagnosis or symptom onset, in all cases, r_s_ = 0.70 (p < 0.0001) (data not shown). Compared to our in-house quantitative RBD ELISA end-point titer data, the sqLFA RI values correlated positively, with an r_s_ of 0.83 (p < 0.0001) (See Supplemental Figure 2B). We then tested the correlation of the DiaSorin Trimeric Spike IgG assay (Liaison, DiaSorin) which was done in a CLIA-certified laboratory. This assay was approved by the Food and Drug Administration in 2021 as “Acceptable for Use in the Manufacture of COVID-19 Convalescent Plasma with High Titers of Anti-SARS-CoV-2 Antibodies” at a cutoff of >/= 87 AU/mL[12]. We found that among n = 94 samples tested, our sqLFA correlated positively with an r_s_ of 0.59 (p < 0.0001) (See Supplemental Figure 2C). Of note, a correlation of the DiaSorin assay result with our NT50 assay resulted in a Spearman correlation of 0.55, which is positive but lower than sqLFA vs NT50 (See Supplemental Figure 2C).

We also tested 38 convalescent serum samples on different days to assess intra-assay variability and found the Spearman correlation to be r_s_ = 0.94 (p <0.0001), with overall coefficient of variation of 82% (n=38), when broken down: 30% for samples with RI > 500 (n=24) and 197% for samples with RI < 500 (n = 14) (data not shown).

### Sensitivity and Specificity Analyses

Multiple ROC analyses were done to calculate the sensitivity and specificity of the sqLFA RI to detect samples with different levels of functional neutralizing antibodies. Samples that had detectable neutralizing antibody titers (NT50 ≥1:20 or other cutoffs) were set as positive controls, and those with undetectable neutralizing antibody titers (or below the cutoff) were set as negative controls using data from our in-house NT50 assay. The area under the curve (AUC) for the analysis to detect any neutralizing antibody NT50 ≥1:20 was 90%, p <0.0001. We found that the sqLFA RI was highly sensitive at the cutoff of >75.5 (∼95%), close to the manufacturer’s cutoff for a positive value (>80), for detecting samples with NT50 ≥1:20 (Table 1). The specificity however, at the cutoff of >75.5 was low, at about 57%. We then looked at higher sqLFA cutoffs in order to find a cutoff that had a high specificity for detecting samples with NT50 ≥1:20. We found that a cutoff of >992 in the sqLFA RI, had a low sensitivity of 40%, but a very high specificity at ∼97% (Table 1). We repeated this analysis for detection of samples with higher NT50s, in order to see what an ideal cutoff would be in order to detect individuals with higher levels of neutralizing antibodies, and found that a highly sensitive cutoff was >448, and a highly specific cutoff of was >1506 (Table 1); the AUC for these further analyses respectively was 85%, 82%, 75%, p<0.0001.

**Table 1.**
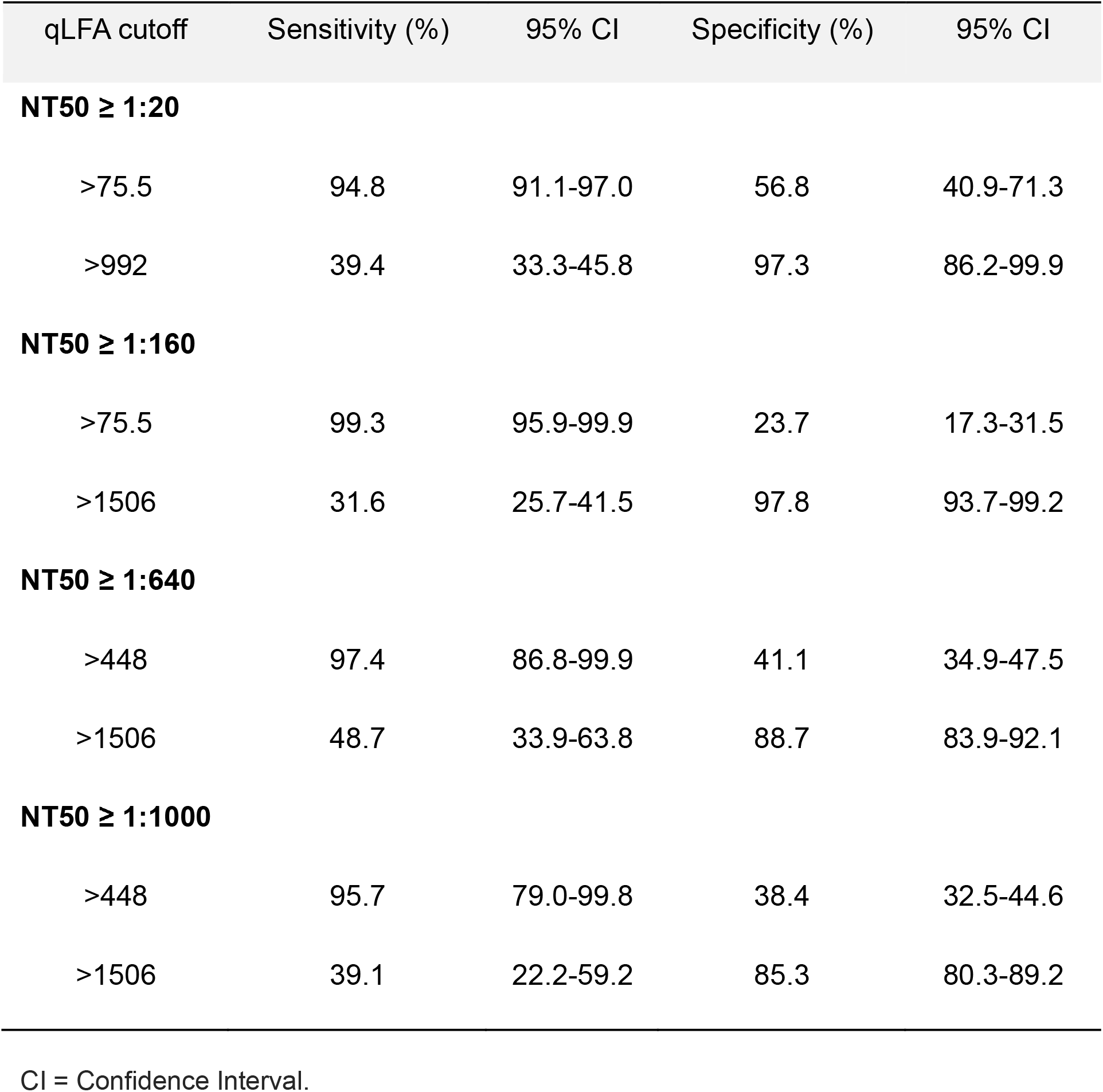
Sensitivity and Specificity of qLFA for detecting neutralizing antibodies.

| qLFA cutoff | Sensitivity (%) | 95% CI | Specificity (%) | 95% CI |
| --- | --- | --- | --- | --- |
| **NT50 ≥ 1:20** |  |  |  |  |
| >75.5 | 94.8 | 91.1-97.0 | 56.8 | 40.9-71.3 |
| >992 | 39.4 | 33.3-45.8 | 97.3 | 86.2-99.9 |
| **NT50 ≥ 1:160** |  |  |  |  |
| >75.5 | 99.3 | 95.9-99.9 | 23.7 | 17.3-31.5 |
| >1506 | 31.6 | 25.7-41.5 | 97.8 | 93.7-99.2 |
| **NT50 ≥ 1:640** |  |  |  |  |
| >448 | 97.4 | 86.8-99.9 | 41.1 | 34.9-47.5 |
| >1506 | 48.7 | 33.9-63.8 | 88.7 | 83.9-92.1 |
| **NT50 ≥ 1:1000** |  |  |  |  |
| >448 | 95.7 | 79.0-99.8 | 38.4 | 32.5-44.6 |
| >1506 | 39.1 | 22.2-59.2 | 85.3 | 80.3-89.2 |

CI = Confidence Interval.

## Discussion

The BioMedomics sqLFA assay can be used as a highly specific tool for the detection of neutralizing antibodies in human plasma or sera and can be adapted to use RBD antigens from different SARS-CoV-2 variants. The sqLFA semi-quantitative readout has a strong positive correlation with NT50 neutralizing antibody titers, and at a cutoff of >992, specifically detects sera positive for functional neutralizing antibodies. This cutoff can be used to identify individuals who may benefit from additional vaccination boosters as well as SARS-CoV-2 exposed or infected individuals who may benefit from monoclonal antibody-based therapies. Furthermore, the even higher cutoff of >1506 on the sqLFA was very specific for individuals with a functional neutralizing antibody titer of NT50 ≥ 1:640.

This new sqLFA can also be used as a highly sensitive screening tool at the manufacturer’s lower cutoff of 80 to identify individuals with any detectable SARS-CoV-2 neutralizing serum antibodies. The low cutoffs we have identified for high sensitivity do however have low specificities, which is common for most screening tools, but indicates the need for a follow up test with higher specificity to eliminate false positives. Furthermore, the sqLFA reader has a small footprint, it is very easy to use, has low intra-assay variability and takes less than 20 minutes to set up and obtain a result, making it a great candidate for clinical use.

Three other rapid, quantitative assays have been published and found to positively correlate with neutralizing antibody levels[13-15]. Two of these assays leverage the interaction between RBD and angiotensin converting enzyme 2 in their lateral flow development. The COVID-19 Nab-test^™^ was studied in a cohort of 79 SARS-CoV-2 infected individuals and found to be correlated with a microneutralization assay (NT50 ≥1:40), R^2^ = 0.72 (p<0.0001)[13]. The quantitative LFA described by Lake *et al*. was compared to neutralization assays from individuals infected and vaccinated and found to have an ROC AUC of 98%[14]. These assays based on RBD antigens, similarly to our assay, show good correlations with neutralizing antibody titers and are a promising start in the development of a rapid quantitative assay surrogate for neutralizing antibody levels and therefore humoral protection against COVID-19 disease.

The main limitation of the sqLFA analyzed here is that the correlation with the sqLFA RI value and NT50 titers is not strong enough to develop specific RI cutoffs in order to group individuals into high versus low neutralizing antibody titer groups with *both high sensitivity and specificity*, for example. However, this is an area of new development and may be improved by using an LFA that simultaneously detects IgM, IgA and IgG in the same strip. Neutralizing antibody assays may have contributions from these other immunoglobulin isotypes especially in the first few months post infection, and this may be an important aspect in the development of a rapid quantitative assay with a stronger positive correlation with neutralization assays. Another limitation is that this sqLFA only detects antibodies against RBD, and though >90% of neutralizing antibodies are directed against RBD[13], some are directed at other areas of the virus that would be missed here. Finally, definitive validation of this assay and reader prototype requires further studies in various populations of individuals infected with different SARS-CoV-2 variants as well as those vaccinated by the available COVID-19 vaccines.

Given ongoing transmission of SARS-CoV-2 variants, as well as known waning antibody levels to both natural infection and vaccination[16], the development of rapid quantitative assays to identify individuals at risk for re-infection is critical. In the SARS-CoV-2 mRNA vaccinated healthcare worker study done by Bergwerk *et al*., individuals in the cohort with breakthrough infections had a 6-7 fold lower mean neutralizing antibody titer than those who had not experienced a vaccine breakthrough infection[1]. Most rapid and non-rapid SARS-CoV-2 serologic assays are developed without direct comparison to functional antibody assays, though it is clear that neutralizing antibodies are an immune correlate of protection against COVID-19. Furthermore, it has been shown that standard two-dose[17] or even three doses[18] of mRNA vaccination in solid organ transplant recipients may not produce an adequate immune response. Current CDC guidelines recommend a three-dose primary schedule followed by a booster at least three months later[19]. Having a reliable, easy to use, rapid clinical assay that detects neutralizing antibody presence and identifies individuals that may need additional SARS-CoV-2 vaccination is an important component of the future management of the ongoing COVID-19 pandemic.

## Abbreviations

sqLFA: Semi-quantitative lateral flow assay
SARS-CoV-2: severe acute respiratory syndrome coronavirus 2
RBD: receptor binding domain
RI: reflective intensity
ELISA: enzyme-linked immunosorbent assay
NT50: 50% neutralization of viral infection
ROC: receiver operating characteristics curve
r_s_: Spearman correlation coefficient
p: p values
LOD: titers below limit of detection
AUC: area under the curve
IgG: Immunoglobulin G

## Acknowledgements

The NIH SeroNet Serocenter of Excellence Award, U54 CA260543, supported generation of laboratory data and the following investigators: AJM, DRB, YJH, RSB, AMD and LAB.

