## Supplemental Figures for "A semi-quantitative, rapid, point of care SARS-CoV-2 serologic assay predicts neutralizing antibody levels"

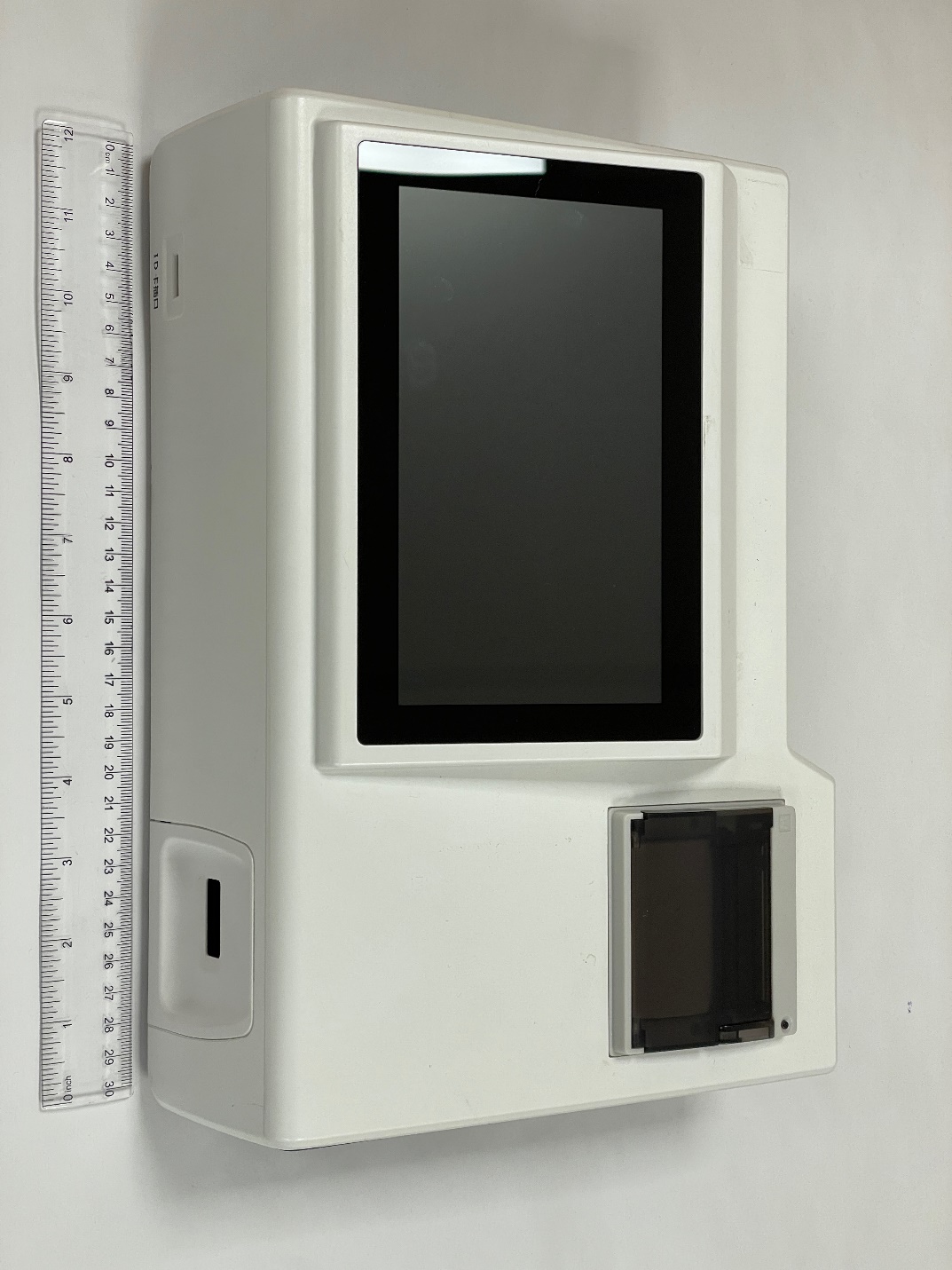

**Supplemental Figure 1**. Image and dimensions of the prototype reader. It is approximately 23 x 30 centimeters (width x length).

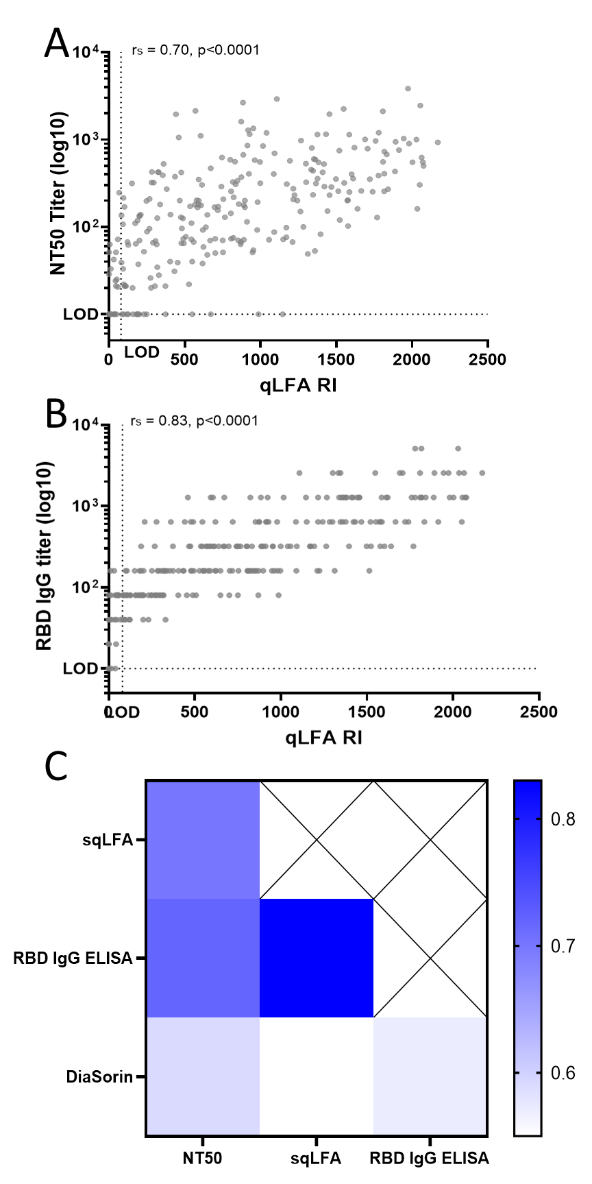

**Supplemental Figure 2**. **qLFA correlation plots**. **A**. sqLFA RI vs NT50 Titer **B**. sqLFA RI vs RBD IgG ELISA end-point titer. **C**. Spearman correlation values for all assays compared, p , 0.0001 for all. For A-C, a non-parametric, two-tailed Spearman’s rank correlation is used to calculate correlation coefficients (r_s_) and p values (p), titers below limit of detection (LOD) for the ELISA and NT50 were set to 10.
